# Machine Learning Predicts Treatment Response in Bipolar & Major Depressive Disorders

**DOI:** 10.1101/2022.05.13.491903

**Authors:** Mustafa S. Salman, Eric Verner, H. Jeremy Bockholt, Zening Fu, Maria Misiura, Bradley T. Baker, Elizabeth Osuch, Jing Sui, Vince D. Calhoun

**Affiliations:** Tri-Institutional Center for Translational Research in Neuroimaging and Data Science (TReNDS) [Georgia State University, Georgia Institute of Technology, and Emory University], Atlanta, GA, USA; School of Electrical & Computer Engineering, Georgia Institute of Technology, Atlanta, GA, USA; Lawson Health Research Institute, London Health Sciences Centre, FEMAP, London, Ontario, Canada; Schulich School of Medicine and Dentistry, Western University, London, Ontario, Canada; Institute of Automation, Chinese Academy of Sciences, and the University of Chinese Academy of Sciences, Beijing, China

**Keywords:** bipolar disorder, major depressive disorder, kernel SVM, medication-class, treatment response, neuromark

## Abstract

Diagnosis of bipolar disorder (BD) patients with complex symptoms presents a challenge to clinicians. Patients tend to spend more time in a depressive state than a manic state. In such complex cases, the current Diagnostic and Statistical Manual (DSM), which is not based on pathophysiology, can lead to misdiagnosis as major depressive disorder (MDD) and an imperfect or even harmful medication response. A biologically-based classification algorithm is needed to improve the accuracy of diagnosis. Osuch et al. (2018) presented a kernel support vector machine (SVM) algorithm to predict the medication-class of response from new patient samples whose diagnoses were unclear. Here we also utilize the kernel support vector machine (SVM) algorithm but with a few novel contributions. We applied the robust, fully automated neuromark independent component analysis (ICA) framework to extract comparable features in a multi-dataset setting and learn a kernel function for support vector machine (SVM) on multiple feature subspaces. The neuromark framework successfully replicates the prior result with 95.45% accuracy (sensitivity 90.24%, specificity 92.3%). To further evaluate the generalizability of our approach, we incorporated two additional datasets comprising bipolar disorder (BD) and major depressive disorder (MDD) patients. We validated the trained algorithm on these datasets, resulting in a testing accuracy of up to 89% (sensitivity 0.88, specificity 0.89) without using site or scanner harmonization techniques. We also translated the model to predict improvement scores of major depressive disorder (MDD) with up to 70% accuracy. This approach reveals some salient biological markers of medication-class of response within mood disorders.

**Highlights:**

- We demonstrate a DSM-free approach for predicting treatment response from resting-state functional magnetic resonance imaging (fMRI) data.
- We identify several replicable biomarkers using the approach.
- Our work has potential for clinical application by replacing trial-and-error in treating complex psychiatric disorders.

## 1. Introduction

Patients without any evident symptoms of mania pose a difficult challenge in differentiating bipolar disorder (BD) from major depressive disorder (MDD) using the current “gold standard,” the DSM (American Psychiatric Association, 2013). BD can, in some cases, be misdiagnosed as MDD because patients with BD generally spend more time in depressive than in manic states (Judd et al., 2002). When it comes to the treatment response of patients to the different medication classes, mood stabilizers (MSs) often do not effectively treat MDD. At the same time, antidepressants (ADs) may worsen BD type I. Fundamental differences between MDD and BD have been widely reported in many cases (Osuch et al., 2018; de Almeida and Phillips, 2013; Bowden, 2005; Perlis et al., 2006). It is necessary to obtain the correct mood diagnosis and medication class to best support the patient’s recovery.

Group independent component analysis (ICA) is a widely used data-driven algorithm for conducting multi-subject fMRI studies. The most widely used spatial group ICA approach identifies maximally spatially independent spatial maps (SMs) using the temporal concatenation of multi-subject data. It subsequently decomposes each subject’s data into unique time courses (TCs) and considerably variable SMs using a back-reconstruction technique (Erhardt et al., 2011). This entirely data-driven approach can be challenging to implement on asynchronous multi-dataset analyses due to the need to analyze all the data together and the complexities of selecting and labeling components. As an alternative, the subject-specific features can be computed using spatially constrained ICA (scICA), which automatically and adaptively estimates individual-level independent components (ICs) using a priori network templates as guidance. Several ICA algorithms available in Group ICA of fMRI Toolbox (GIFT) (https://trendscenter.org/software/gift/) can be used for scICA (Lin et al., 2010; Du and Fan, 2013). This work combines scICA with component templates derived and replicated from multiple large N (N>800) data sets and takes 74 individual subject’s fMRI data as input.

Support vector machines (SVMs) are a set of supervised binary classification algorithms which can also be extended for regression and multiclass classification (Vapnik, 1999, 1998). The SVM algorithm incorporates a sample selection mechanism, i.e., only the support vectors affect the decision function. It constructs a maximal margin linear classifier in high-dimensional feature space by mapping the original features via a kernel function. We can define a unique kernel function for applying the SVM algorithm to classify fMRI features such as SMs and TCs. We utilize the similarity measures of subspaces in the commonly used kernel functions (Chang and Lin, 2001).

Here we extended the classification algorithm proposed in prior work to multi-feature (i.e., using SMs and TCs) and multi-dataset cases (Osuch et al., 2018; Fan et al., 2011). We used BD and MDD subjects from three different datasets (Western, UCLA Consortium for Neuropsychiatric Phenomics LA5c (LA5C), and Establishing Moderators and Biosignatures of Antidepressant Response for Clinical Care for Depression (EMBARC)) and controls to create/validate a new SVM-based classification algorithm (Osuch et al., 2018; Poldrack et al., 2016; Trivedi et al., 2016). We also used feature selection on the model trained using known DSM-based MDD vs. BD, type I patients to reveal neurophysiological differences in these populations. The longer-term hope is that the algorithm will be helpful to predict AD vs. MS response in complex patients whose DSM diagnoses are unclear. The focus on medication class response provides a ’DSM-free’ and potentially clinically useful approach to identifying biological markers of medication-class of response within mood disorders.

This work extends our prior work in several ways (Salman et al., 2021). Previously, we reported results using thresholded SMs as features. Here we use the unthresholded SMs, which lowers the number of false positives and false negatives and results in better sensitivity/specificity. We also use the unthresholded SM in conjunction with functional network connectivity (FNC) to perform multi-feature prediction in Western data and extend the framework in LA5C and EMBARC data. We also include the prediction of MDD patient treatment response improvement scores in the EMBARC dataset using the same algorithm.

## 2. Methods

### 2.1. Data

Our medication-class of response predictor model is trained on the resting-state fMRI data collected on MDD and BD patients from Western University. These individuals were followed up over an extended time period and categorized based on the cumulative knowledge including medication class response as well as both clinical and research diagnosis. We validated the trained model on two independent datasets: EMBARC and LA5C. These datasets are described below and also summarized in Tab. 1.

**Table 1:** Summary of data

| **Dataset** | **Western** | **LA5C** | **EMBARC** |
| --- | --- | --- | --- |
| **Population info** |  |  |  |
| **BD** | 44 | 49 |  |
| **MDD** | 43 |  | 545 |
| **Nonresponder** | 14 |  |  |
| **Control** | 39 | 121 | 78 |
| **Total** | 140 | 170 | 623 |
| **Age range** | 16-27 | 21-50 | 18-65 |
| **Acquisition parameters** |  |  |  |
| **Scanner type** | Siemens | Siemens | GE/Siemens/Phillips |
| **TR** | 2000 ms | 2000 ms | 2000 ms |
| **TE** | 30 ms | 30 ms | 28 ms |
| **Slices** | 40 | 34 |  |
| **Slice thickness** | 3 mm | 4 mm | 3.1 mm |
| **Flip angle** | 90° | 90° | 90° |
| **FOV** | 240 mm | 192 mm | 205 mm |
| **Matrix size** | 80x80 | 64x64 | 64x64 |
| **Scan duration** | 8 minutes | 304 s | 2x6 minutes |
| **Volumes** | 164 | 142 | 178 (-4) |

#### 2.1.1. Western Data

The University of Western Ontario Research Ethics Board approved the data collection. All participants, aged 16-27, provided written informed consent. A total of 140 subjects were divided into four groups as follows: 33 controls, 35 MDD, 35 BD, 12 unknown diagnosis, and 25 scanned depressed before being treated. Additional details about the participants and data acquisition can be found in prior work (Osuch et al., 2018).

MRI data were acquired using a 3.0T Siemens Verio MRI scanner at the Lawson Health Research Institute using a 32-channel phased-array head coil. fMRI data included gradient-echo, echo-planar imaging (EPI) scans with repetition time (TR) = 2000 ms, echo time (TE) = 30 ms, 40 axial slices and thickness = 3*mm,* with no parallel acceleration, flip angle = 90°, field of view (FOV) = 240 × 240mm, matrix size = 80 × 80. The resting fMRI scan lasted approximately 8 minutes (164 brain volumes). After preprocessing and quality control, 135 subjects were retained: 13 non-responders or nonresponder (remitted without medication), 41 responding to AD, 42 responding to MS, and 39 controls.

#### 2.1.2. LA5C Data

The data included imaging data from healthy individuals from the community (130 subjects) and samples of individuals diagnosed with schizophrenia (50), bipolar disorder (49), and attention deficit hyperactivity disorder (ADHD) (43). MRI images were collected on two 3T Siemens Trio scanners located at the Ahmanson-Lovelace Brain Mapping Center (Siemens version syngo MR B15) and the Staglin Center for Cognitive Neuroscience (Siemens version syngo MR B17) at UCLA.

Functional scans were acquired using a T2*-weighted EPI sequence with the following parameters: TR = 2000 ms, TE = 30 ms, slice thickness = 4 mm, 34 slices, oblique slice orientation, flip angle = 90°, matrix 64 × 64, and FOV = 192 mm. Scans covered the whole brain for a total time of 304 s. After preprocessing and quality control, 255 subjects were retained: 121 controls, 46 diagnosed with BD, 47 schizophrenia (SZ), and 41 ADHD. For this analysis, we used the controls and BD subjects. Additional data descriptions can be found in prior work (Poldrack et al., 2016).

#### 2.1.3. EMBARC Data

Functional imaging was acquired during the resting-state for 2 scans of length 6 minutes each. Outpatients aged 18-65 with MDD, diagnosed using the Structured Clinical Interview for DSM-IV Axis I Disorders (SCID), were recruited at four sites. The functional image acquisition parameters were: TR = 2000 ms, TE = 28 ms, flip angle = 90°, FOV = 205 mm, slice thickness = 3.1 mm, matrix 64 × 64. After preprocessing and quality control 623 subjects were retained: 78 controls, and 545 diagnosed with MDD. Additional details about the data acquisition can be found in prior work (Trivedi et al., 2016).

The patients were divided into four groups between 1 and 4 based on the treatment response. Response to treatment was defined by being at least “much improved” (3) on the Clinical Global Improvement (CGI) scale for patients (Trivedi et al., 2016).

### 2.2. Preprocessing

We preprocessed all subjects’ data from the three different datasets using the Statistical Parametric Mapping (SPM) software (Friston, 2007). We performed rigid body motion correction to correct for subject head motion, followed by the slice-timing correction to account for the timing difference in slice acquisition. We subsequently warped the fMRI data into the standard Montreal Neurological Institute (MNI) space using an EPI template and resampled the data to 3 × 3 × 3 *mm*^3^ isotropic voxels. We further smoothed the resampled fMRI images using a Gaussian kernel with a full width at half maximum (FWHM) of 6 mm.

We took the following quality control measures after preprocessing. We retained a subject for analysis if they had: (1) data with head motion amounting to less than 3° rotation and 3mm transition along the whole scanning period; (2) data with more than 120 time points in the fMRI acquisition; (3) data providing a successful normalization to the whole brain as assessed by visual inspection and spatial correlation with the template (Fu et al., 2021b).

### 2.3. Feature Extraction

#### 2.3.1. The neuromark Template

We used a labeled and ordered set of IC templates called neuromark (https://trendscenter.org/data/) (Du et al., 2020). This reference set comprised 53 components which replicated following separate analyses of two independent datasets consisting of controls: Human Connectome Project (HCP) and Brain Genomics Superstruct Project (GSP) (Smith et al., 2013; Buckner et al., 2014). The components were divided into seven functional domains: subcortical network (SCN), auditory network (ADN), sensorimotor network (SMN), visual network (VSN), executive control network (CON), default mode network (DMN), and cerebellar network (CBN). Fig. 2 presents a composite view of the neuromark templates. The neuromark templates have been successfully applied in numerous studies and have been validated as robust spatial priors that provide reliable functional network features across subjects and datasets (Fu et al., 2021c,b,a).

#### 2.3.2. Spatially-constrained ICA

We used the spatially-constrained ICA algorithm available in GIFT (https://trendscenter.org/software/gift/ to extract features from preprocessed data of the subjects (Du and Fan, 2013; Lin et al., 2010; Salman et al., 2019). We used the neuromark network templates and each subject’s fMRI data as inputs to derive subject-specific SMs and TCs in a fully automated approach. We then calculated the Pearson correlation coefficient between the TCs from 53 neuromark components of each subject and obtained a FNC matrix with dimension 53 × 53.

#### 2.3.3. Kernel SVM

Fig. 1 presents a flowchart of the classification framework. Given two sub-spaces **A** = {**a**_1_, **a**_2_,…, **a**_*k*_,} and **B** = {**b**_1_, **b**_2_,…, **b**_*k*_,}, where {**a**_1_, **a**_2_,…, **a**_*k*_, } and {**b**_1_, **b**_2_,…, **b**_*k*_, } are orthonormal basis vectors (i.e., the component spatial maps) for subspaces **A** and **B**, the subspace distance metric can be obtained from the ordered singular values of **A**’**B**, i.e., 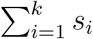, where *s* = *svd*(**A**′**B**) (Björck and Golub, 1973; Fan et al., 2011).

**Figure 1:**
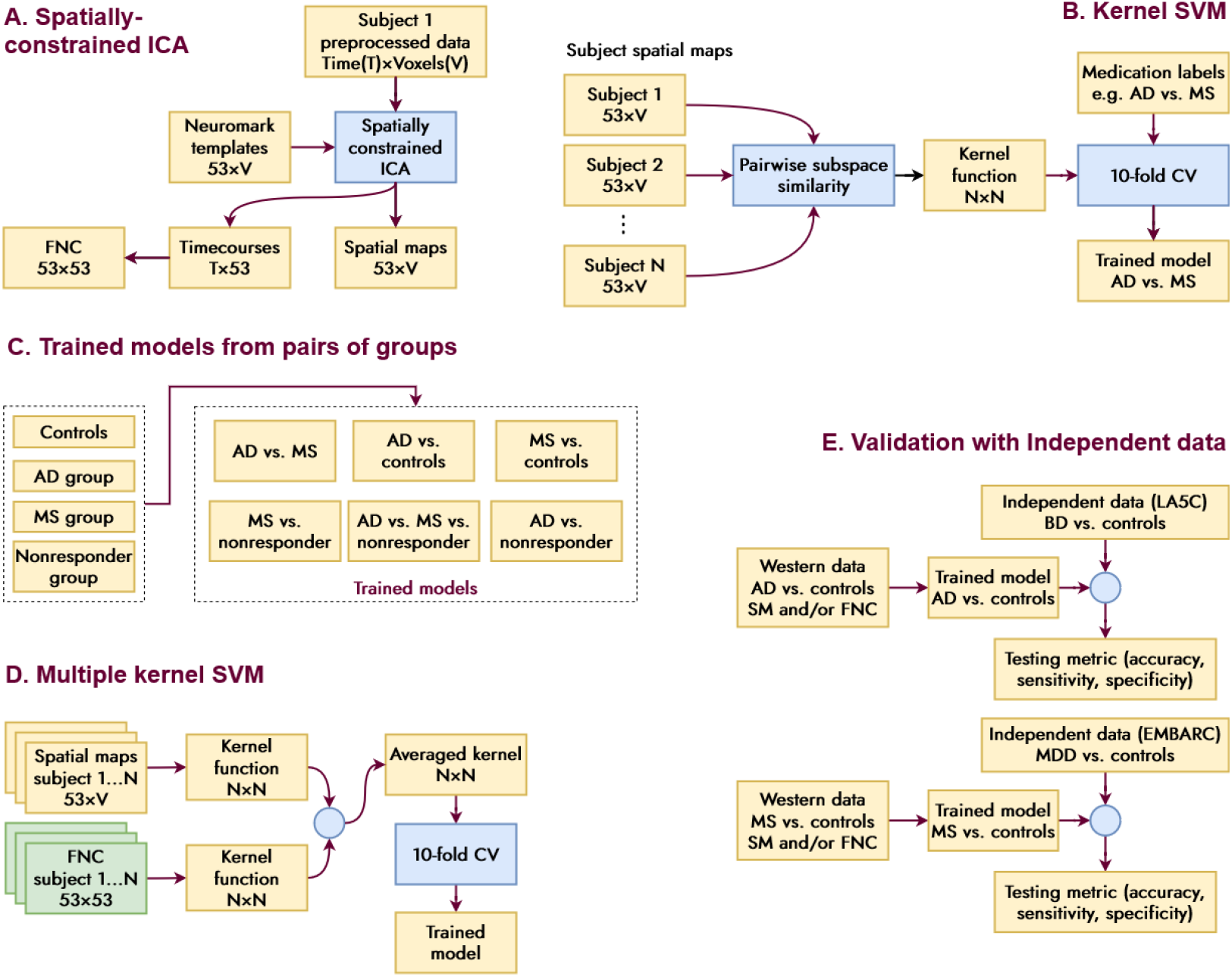
Flowchart of our classification scheme. A. Resting-state fMRI data are put through the Neuromark ICA pipeline for feature (spatial maps, time courses and FNC) extraction. B. Classification is performed using Kernel SVM algorithm & 10-fold crossvalidation. C. Known medication-class of treatment response (mood stabilizers (MS)/anti-depressants (AD) and controls/nonresponders) are used as the targets to train the models. D. Experiments are run using spatial maps (SMs), functional network connectivity (FNC) and their combination as features. E. Trained models are tested on independent data.

**Figure 2:**
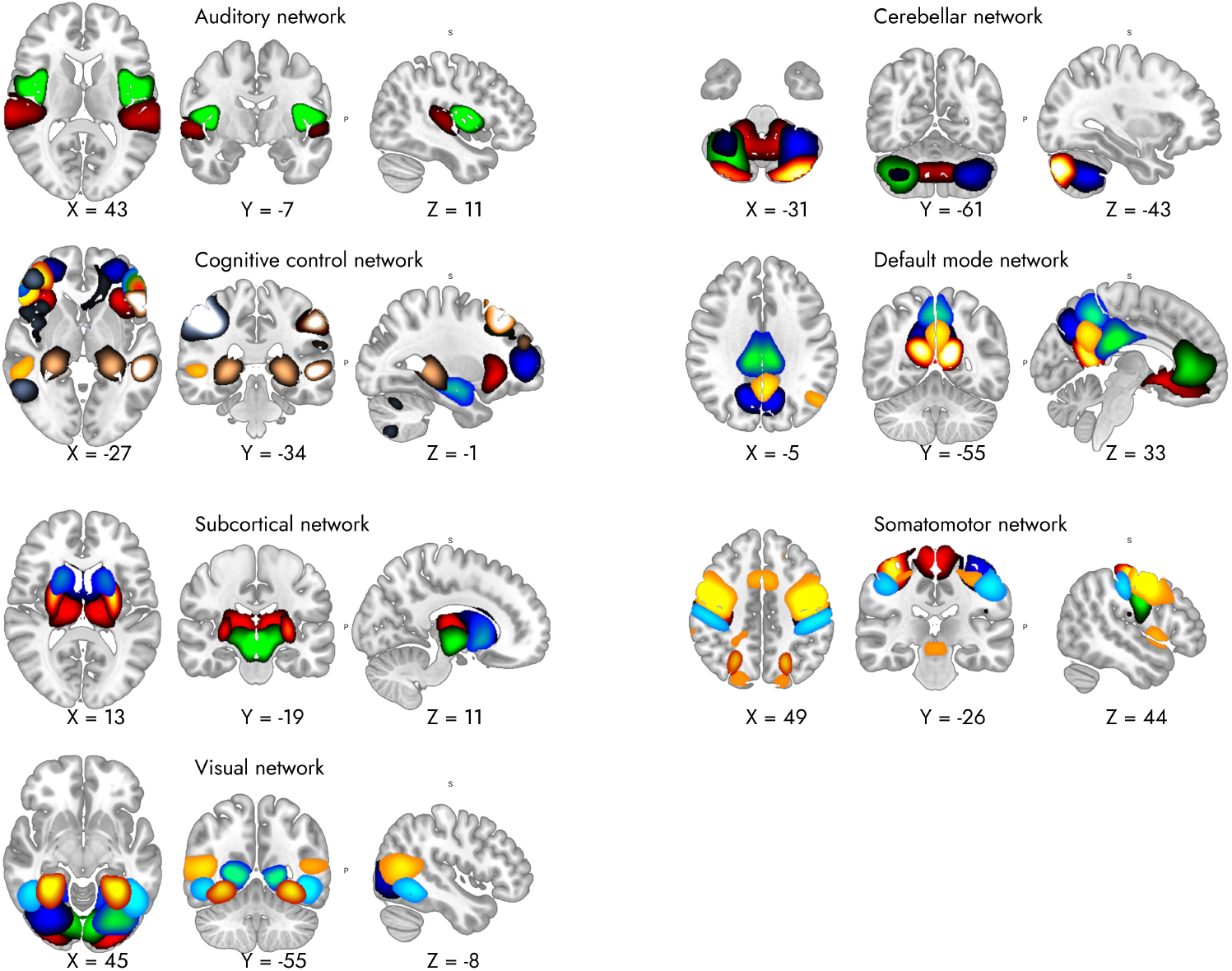
The neuromark SM templates. These are obtained using group ICA analysis on HCP and GSP controls data and a greedy algorithm for identifying the most replicable components. 53 spatial maps are divided into 7 functional domains. These templates can be used as references to estimate subject spatial maps and time courses from new and unseen data.

Then subspace similarity between different subjects is defined correspondingly as

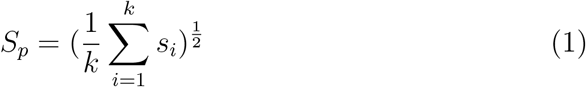

Here *p* is short for projection, and *k* is the number of subspace dimensions. Suppose the number of BD and MDD are *m* and *n*, respectively; we can construct an (*m*+*n*)-dimensional symmetric matrix based on the Riemannian similarity metric between different subjects as in Eq. 1. This derivation is possible due to the (maximally) statistically independent nature of the subject-specific SMs, which approximate an orthonormal set.

Next, *S_p_* is mapped into a high-dimensional feature space via a sigmoid kernel function as defined in

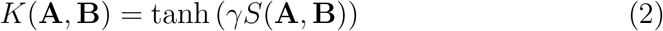

where *S*(**A**, **B**) is the similarity in Eq. 1, and *γ* is the kernel parameter. With this kernel function, we can build a SVM classifier.

We used the following formulation to estimate subspace similarities as we deal with data containing features from multiple modalities (SMs and TCs/FNC). Given two subspaces (**A_1_**, **A_2_**) = ({**a**_11_, **a**_12_,…, **a**_1*k*_}, {**a**_21_, **a**_22_,…, **a**_2*k*_}) and (**B_1_**, **B_2_**) = ({**b**_11_, **b**_12_,…, **b**_1*k*_}, {**b**_21_, **b**_22_,…, **b**_2*k*_}), where **A_1_**, **B_1_** are sub-spaces for the subjects from one modality, and **A_2_**, **B_2_** are subspaces from another modality, the subspace similarity between the two subjects are defined by extending Eq. 1 as

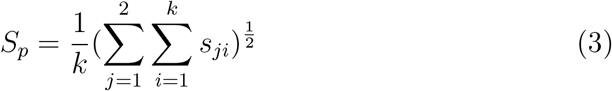

#### 2.3.4. Forward Feature Selection

We used a stepwise forward selection method to generate an optimal component set (Osuch et al., 2018). We used the Matlab sequentialfs function along with 10-fold cross-validation (CV). Algorithm 4 contains the pseudocode for implementing this step.

1. Start with empty set *Y*_0_ = *ϕ* and some candidate features
2. Select the best next feature 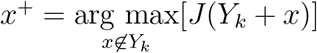
3. Update *Y*_*k*+1_ = *Y_k_* + *x*^+^; *k* = *k* + 1
4. Return to 2

### 2.4. Experiments

We performed each of the following experiments using a 10-fold CV around the kernel SVM framework described above. We repeated each experiment 100 times by shuffling the CV folds to ensure the stability of the reported results.

#### 2.4.1. Classification Between different Groups in Western Data

We first replicated the result reported in Osuch et al. using the neuro-mark ICA framework (Osuch et al., 2018). For this experiment, we used the patients with known medication-class of response (*N* = 66) as used in the prior work. We used 66 patients’ data as the input and their treatment response (MS or AD) classification labels. We performed three different experiments with different features from the same subjects: SMs, FNC in the kernel SVM approach, and combined SM+FNC in the multiple kernel learning approach. In the next experiment, we added more data coming from the same site (*N* = 83) for replication.

#### 2.4.2. Classification of Controls and nonresponder Subjects

Two more cohorts are present in the Western data, namely, remitted without medication (nonresponder) and controls. Therefore, we separately performed the classification of medication-class of response between the nonresponder-MS and nonresponder-AD groups and classification between controls-BD and controls-MDD groups.

### 2.5. Classification of BD and MDD in LA5C and EMBARC Data

For the validation experiments, we used data from LA5C and EMBARC projects. We trained the classification models using the controls-BD cohorts of the Western data and the three sets of features mentioned above and then tested them on the same cohort in the LA5C data. Similarly, for EMBARC, we trained the models using the controls-MDD populations of the Western data and then tested them on the same cohort in the EMBARC data. Note that according to Tab. 1 both LA5C and EMBARC datasets suffer from unbalanced sample issue. To mitigate this problem, we used a subject selection step before classification. In this step, we selected equal number of BD and MDD patient data as the number of controls data from LA5C and EMBARC, respectively. We also classified the patients who responded well (improvement score of 4) and nonresponders (improvement score of 1) in the EMBARC data.

### 2.6. Most Salient Features

Previously we mentioned that we performed a forward feature selection procedure to reduce the 53 components estimated using neuromark ICA into a smaller set of SMs. We performed this step in every CV step of every experiment, which gave us a set of most frequently occurring discriminative SMs. We noted these salient brain activity patterns for discussion.

## 3. Results

### 3.1. Classification Between different Groups in Western Data

Tab. 2 lists the results of the classification experiments. We used subjects from Western data to train the models in all of these experiments. We repeated every experiment 100 times with shuffled CV folds to ensure reproducibility. Below we report average metrics across those runs for each experiment.

**Table 2:** Classification results summary

| Groups | Features | Training N | Testing scores (hold-out data) |  |  |
| --- | --- | --- | --- | --- | --- |
| Osuch et. al. (2018) |  |  | Accuracy | Sensitivity | Specificity |
| MS-AD | SM | 64 | 92.84 ± 1.88 | 0.90 | 0.92 |
| Replication using Neuromark ICA |  |  |  |  |  |
| MS-AD | SM | 83 | 84.33 ± 3.34 | 0.87 ± 0.03 | 0.80 ± 0.05 |
|  | FNC |  | 85.48 ± 2.93 | 0.91 ± 0.01 | 0.79 ± 0.05 |
|  | SM+FNC |  | 86.06 ± 2.59 | 0.90 ± 0.01 | 0.81 ± 0.05 |
| MS-nonresponders | SM | 54 | 93.27 ± 2.89 | 0.99 ± 0.01 | 0.71 ± 0.12 |
|  | FNC |  | 97.09 ± 1.32 | 1.00 | 0.86 ± 0.05 |
|  | SM+FNC |  | 97.11 ± 1.21 | 1.00 | 0.87 ± 0.05 |
| AD-nonresponders | SM | 53 | 92.38 ± 2.34 | 0.99 ± 0.01 | 0.68 ± 0.11 |
|  | FNC |  | 97.33 ± 0.93 | 1.00 | 0.88 ± 0.04 |
|  | SM+FNC |  | 97.47 ± 1.11 | 1.00 | 0.88 ± 0.04 |
| MS-AD-nonresponders | SM | 95 | 80.24 ± 2.85 | 0.83 ± 0.02 | 0.81 ± 0.07 |
|  | FNC |  | 88.18 ± 2.52 | 0.87 ± 0.01 | 0.89 ± 0.05 |
|  | SM+FNC |  | 87.33 ± 2.63 | 0.86 ± 0.01 | 0.87 ± 0.05 |
| MDD-NC | SM | 80 | 85.16 ± 3.02 | 0.84 ± 0.04 | 0.85 ± 0.03 |
|  | FNC |  | 84.49 ± 2.61 | 0.90 ± 0.02 | 0.78 ± 0.05 |
|  | SM+FNC |  | 84.41 ± 2.52 | 0.90 ± 0.02 | 0.78 ± 0.05 |
| MDD-NC (EMBARC) | SM | 77 | 89.00 ± 3.74 | 0.88 ± 0.02 | 0.89 ± 0.06 |
|  | FNC |  | 87.62 | 0 | 1 |
|  | SM+FNC |  | 85.84 ± 3.39 | 0.90 ± 0.01 | 0.81 ± 0.06 |
| BD-NC | SM | 81 | 85.88 ± 3.41 | 0.85 ± 0.04 | 0.85 ± 0.04 |
|  | FNC |  | 85.70 ± 2.59 | 0.91 ± 0.01 | 0.79 ± 0.04 |
|  | SM+FNC |  | 85.17 ± 2.49 | 0.90 ± 0.02 | 0.79 ± 0.05 |
| BD-NC (LA5C) | SM | 166 | 79.75 ± 1.82 | 0.50 ± 0.06 | 0.90 ± 0.02 |
|  | FNC |  | 82.59 ± 0.76 | 0.63 ± 0.01 | 0.89 ± 0.01 |
|  | SM+FNC |  | 82.54 ± 0.84 | 0.63 ± 0.02 | 0.89 ± 0.01 |

The first row is a reproduction of the result reported by Osuch et al. (Osuch et al., 2018) using the SM features. We used neuromark ICA-generated features from 66 MS and AD responder subjects to train the SVM model. The hold-out testing accuracy, in this case, was 92.84%. There were 12 subjects with unknown diagnoses in the original paper. The model achieved 90.9% accuracy in classifying those ”unknown” samples. When classifying newly obtained data from the same site (*N* = 83), the accuracy was reduced to 84.33% (sensitivity 0.87, specificity 0.80).

When similarly classifying nonresponder subjects from MS we obtained 93.27% accuracy (sensitivity 0.99, specificity 0.71, *N* = 54). The same classification approach with AD resulted in 92.38% accuracy (sensitivity 0.99, specificity 0.68, *N* = 53). In a 3-way classification among MS, AD responders and the nonresponder subjects (*N* = 95), we obtained 80.24% accuracy (sensitivity 0.83, specificity 0.81). In most of these experiments, the sensitivity and specificity were quite high, as reported in the table, and generally, sensitivity was slightly higher than specificity.

We also report FNC-based classification results in each alternate row of Tab. 2. We see that the FNC outperformed the SM in classifying these cohorts in all experiments.

### 3.2. Classification in Independent Data

#### 3.2.1. Classification of BD in LA5C data

In the validation experiment using LA5C data consisting of controls and BD populations, the model using SM features achieved a hold-out testing accuracy of 85.88% (sensitivity 0.85, specificity 0.85). The accuracy was 85.88% for BD type-1 diagnosed patients in Western data.

#### 3.2.2. Classification of MDD in EMBARC data

In the other validation experiment using EMBARC data consisting of controls and MDD populations, the model achieved a hold-out testing accuracy of 87.48% (sensitivity 0.85, specificity 0.84). The accuracy was 85.16% for BD-diagnosed patients in Western data.

##### Classification Based on MDD Patient Improvement Scores in EMBARC data

The algorithm was able to separate patients with an improvement score of 1 (no improvement) and 4 (completely improved) with an accuracy of 69.44% (sensitivity 0.96, specificity 0.09).

### 3.3. Most Salient Features

Fig. 3 shows multi-planar views of the neuromark templates whose corresponding subject-level features were the best-performing ones across all the experiments. The Automated Anatomical Labeling (AAL) labels for these templates are the following: superior temporal gyrus from the ADN network (volume 21 in the neuromark template), cerebellum from the CBN network (13), inferior parietal lobule from the CON network (68), precuneus from the DMN network (32), caudate from the SCN network (69), postcentral gyrus from the SMN network (3), and calcarine gyrus from the VSN network (16).

**Figure 3:**
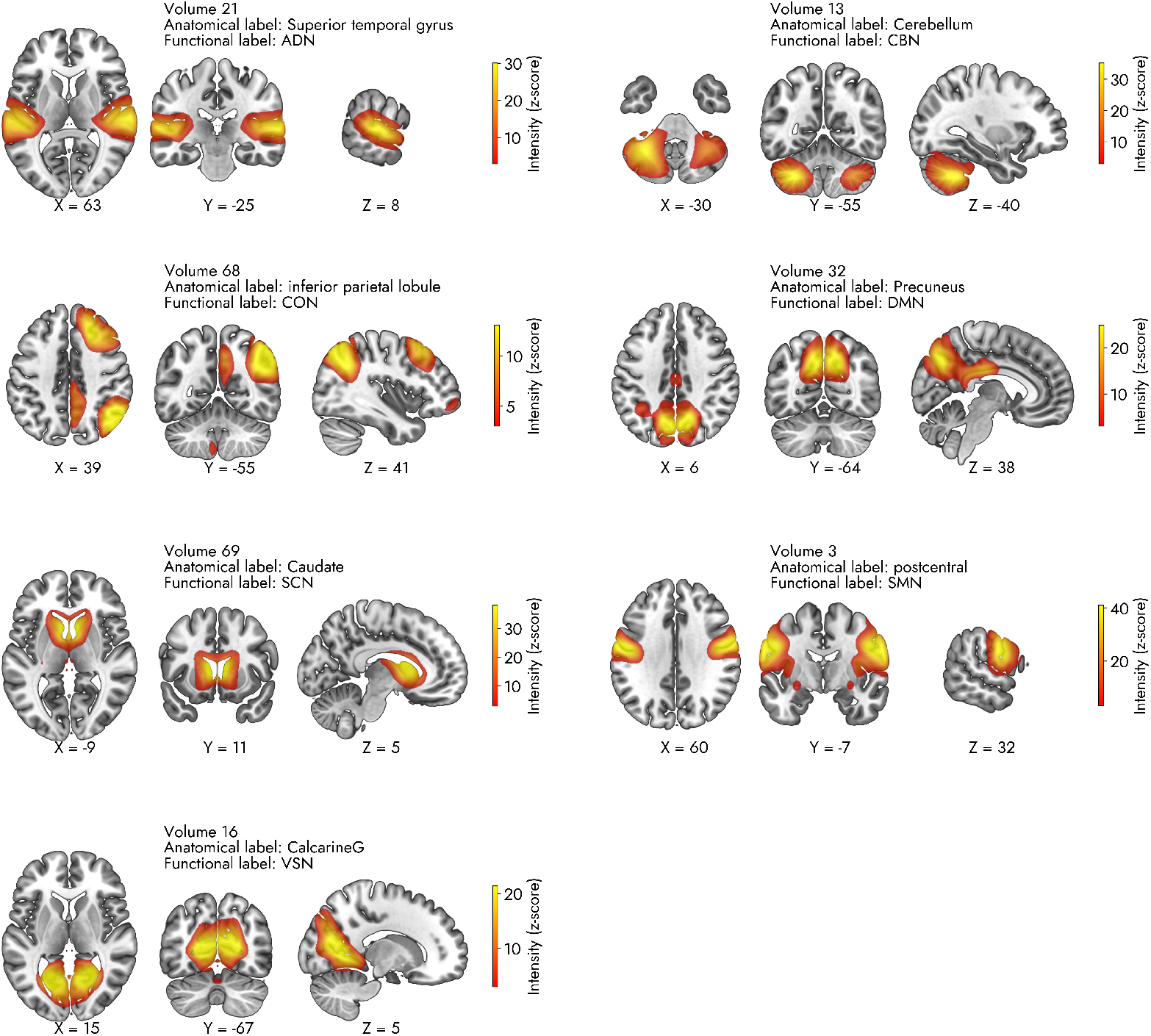
Best component(s) in each functional domain across different experiments

## 4. Discussion

We validated the models trained on the SM and FNC features of the subjects in the Western data by testing them on the LA5C and EMBARC data. The experiments using the FNC features performed better compared to SM. There are several potential factors affecting the performance. There is an age difference across the datasets. The treatment response information is available from Western data only; the other sites provide DSM diagnosis. The datasets are imbalanced, as there are more BD and MDD samples in LA5C and EMBARC datasets than controls. Measures such as regressing the effect of site and other variables such as age from the SMs and FNC or implementing multi-site harmonization techniques such as ComBat may further improve results (Johnson et al., 2007; Fortin et al., 2017, 2018).

Regarding the most salient components in classification, there are several areas of agreement between the neuromark templates and prior work (Osuch et al., 2018). Namely, the precuneus in DMN, postcentral gyrus in SMN, and superior temporal gyrus in ADN appear to be discriminative in both. One notable difference is the model order of ICA; in prior work, the model was set to 20 to reduce the computational cost, whereas the neuromark networks are selected from a group ICA analysis of model order 100.

We also separately ran classification experiment on the independent datasets using 10-fold CV and SM and FNC features. In addition, we used the medication-class of response as the target label in the Western data but the diagnosis label in the case of LA5C and EMBARC data. In doing so, we can demonstrate that the model trained on treatment response data can also predict the DSM diagnosis, although perhaps slightly less accurately. Another versatile feature of the model is the prediction of MDD improvement scores with reasonable accuracy. The low specificity of most of these experiments indicates comparatively high false positives than false negatives when detecting improvement scores of the patients.

We used Eq. 3 to generate a kernel function consisting of multiple modalities. This process can be improved by using a weighted approach or multiple kernel learning approach (Tanabe et al., 2008; Gönen and Alpaydin, 2011). The Riemannian distance measure is most useful when orthonormal basis vectors span the subspaces. Here, one of the modalities in question (FNC) is not spanned by vectors but consists of scalar values. The nature of spatially constrained ICA or spatial group ICA algorithms dictates that the SMs are statistically independent but not necessarily the TCs. This method may fare better in the multi-modal analysis if we extract orthonormal features from the TCs.

Another limitation of the study is that the samples from the different cohorts were imbalanced in some experiments. Therefore the accuracy metric may not convey the efficacy of the method entirely. We have addressed this issue by sampling balanced subjects and reporting more meaningful metrics such as sensitivity and specificity. Also, we treat the SMs and FNC as different modalities in the machine learning sense. Still, they are derived from the same modality (fMRI) in the neuroimaging sense. Future work may rely on more modalities such as structural magnetic resonance imaging (sMRI) and diffusion tensor imaging (DTI) data of the subjects to undertake truly multi-modal analysis.

